# Hormetic effect induced by *Beauveria bassiana* in *Myzus persicae*

**DOI:** 10.1101/2024.01.04.574279

**Authors:** Leonhard Satrio Arinanto, Ary Anthony Hoffmann, Perran Albert Ross, Xinyue Gu

## Abstract

*Myzus persicae*, a serious sap-sucking pest of a large variety of host plants in agriculture, is traditionally controlled using chemical insecticides but there is interest in using biopesticides as restrictions are increasingly placed on the use of broad-spectrum pesticides. Here we show that in petri dish experiments high concentrations of the fungal entomopathogen *Beauveria bassiana* (strain PRRI 5336) lead to rapid mortality of *M. persicae* but at a low concentration (1 × 10^4^ conidia mL^−1^) there is a hormetic effect where longevity and fecundity are enhanced. Hormetic effects persisted across a generation with reduced development times and increased fecundity in the offspring of *M. persicae* exposed to *B. bassiana*. Whole plant experiments point to a hormetic effect being detected in two out of three tested lines. The impact of these effects might also depend on whether *M. persicae* was transinfected with the endosymbiont *Rickettsiella viridis*, which decreases fecundity and survival compared to aphids lacking this endosymbiont. This fecundity cost was ameliorated in the generation following exposure to the entomopathogen. While *B. bassiana* is effective in controlling *M. persicae* especially at higher spore concentrations, utilization of this entomopathogen requires careful consideration of hormetic effects at lower spore concentrations, and further research to optimize its application for sustainable agriculture is recommended.

**AUTHOR SUMMARY:** Biopesticides such as *Beauveria bassiana* can be effective alternatives to chemical insecticides to control insect pests. We tested the efficacy of this biopesticide against the important agricultural pest aphid *Myzus persicae* in laboratory experiments. We also tested whether the potential biological control agent and endosymbiont *Rickettsiella viridis* could provide protection against mortality caused by *B. bassiana*. While high doses of *B. bassiana* caused rapid mortality in aphids, low doses enhanced aphid fecundity and survival. This enhancement persisted into the next generation, with shortened development times and increased fecundity regardless, even when high doses were used in the previous generation. The endosymbiont *R. viridis* did not provide clear protection against *B. bassiana*, in contrast to previous studies in other aphid species, but beneficial effects at low doses also occurred in this aphid line. We also observed hormesis on experiments on whole plants, but only for some aphid genotypes. To a lesser extent, we also observed beneficial effects of low doses of *B. bassiana* in experiments on whole plants, but only in some aphid genotypes. Fitness enhancement by biopesticides at low doses raises concerns for field applications but further research is required to understand its underlying mechanisms.

## INTRODUCTION

Aphids (Homoptera: Aphididae) are sap-feeding insects that pose significant threats to both crops and ornamental plants. Worldwide, there are currently more than 5,000 aphid species (Favret 2022), with around 450 species exclusively associated with crops and approximately 100 species having effectively adapted to agricultural environments, causing substantial challenges for the agricultural industry (Blackman and Eastop 2017). In optimal growth conditions, aphids can rapidly reproduce, spread to neighboring plants, and overwhelm a plant’s capacity to host aphids. Some aphids can transmit arboviruses and alter plant physiology via secretion of phytotoxic saliva (Ng and Perry 2004; Ogawa and Miura 2014; Kumar 2020). Moreover, aphids in general can induce indirect damage by promoting the growth of black sooty mold on leaves through the secretion of honeydew (Kumar 2020).

*Myzus persicae* (Sulzer) or green peach-potato aphids stand out as one of the most successful aphid species in agricultural landscapes. Aside from their primary host (plants from the genus *Prunus*), they have adapted to feed on over 400 secondary host plant species, including economically important crops (Blackman and Eastop 2000). *Myzus persicae* transmits more than 100 arboviruses (Van Emden et al. 1969) and has developed resistance to broad range insecticides (Babineau et al. 2020). Consequently, they are one of the most significant pest species globally as they cause substantial financial losses, reduced crop yields, and are increasingly harder to control using chemical insecticides. For instance, Valenzuela and Hoffmann (2015) estimated that direct feeding and virus-associated injuries result in a 14 to 43% reduction in the yield of grain crops, equating to total annual economic losses of $241 million and $482 million associated with feeding and virus injuries, respectively.

Chemical insecticides remain the primary means to control *M. persicae*, owing to their efficiency, availability, and affordability (Aktar et al. 2009). However, the adverse effects linked to chemical insecticides, including harm to non-target beneficial insects, implications for human health and food safety, as well as insecticide resistance development, drives the need for sustainable alternative control strategies (Kingsley-Nwosu, and John, 2022; Nicolopoulou-Stamati et al., 2016; Sawicki and Denholm, 2008). One alternative strategy utilizes entomopathogenic fungi to suppress aphid populations (Bamisile et al. 2021). Briefly, entomopathogenic fungi are soil-dwelling fungi that infect insects by growing spores that penetrate the insect’s cuticle, leading to systemic infection and death. *Beauveria bassiana* (Balsamo) Vuillemin (Ascomycota: Hypocreales) is a generalist entomopathogenic fungi and some strains are optimized to eliminate pest insects including *M. persicae* (Biryol et al. 2022; Ni et al. 2023). Despite its lethality to insects, commercially available *B. bassiana* strains are typically safe for vertebrates and the environment (Zimmermann 2007), making them an appealing alternative to chemical insecticides. However, there are several challenges in using entomopathogenic fungi as a bioinsecticide to control aphids, including the presence of symbiotic intracellular bacteria that are associated with increased aphid survival in the presence of entomopathogenic fungi infections (Łukasik, van Asch, et al. 2013; Scarborough et al. 2005; Ali et al. 2022).

*Rickettsiella viridis* is a native secondary intracellular bacterial symbiont of pea aphids *Acyrthosiphon pisum* (Harris) (Tsuchida et al. 2010, 2014). This symbiont increases the survival of *A. pisum* against entomopathogenic fungi infection (Łukasik, Guo, et al. 2013). In a recent study, *R. viridis* was introduced into *M. persicae*, a novel aphid host, through microinjection, resulting in substantial deleterious effects and rapid spread within the *M. persicae* population (Gu et al. 2023). Symbionts including *R. viridis* have potential biological control applications due to their ability to induce deleterious effects. However, there is currently no study which reports the effects of *R. viridis* in *M. persicae* against fungal pathogens.

This study investigates the fitness effects of *R. viridis*-infected and *R. viridis*-free *M. persicae* against *B. bassiana* at different spore concentrations with different methods. We used a commercially available entomopathogenic fungus strain of *B. bassiana* PRRI 5539, which was first isolated from *Conchyloctenia punctata* by Dr Schalk Schoeman, and has insecticide activity against *M. persicae* (BASF 2014). Contrary to our initial hypothesis, which proposed that *B. bassiana* would adversely affect the survival and fecundity of *M. persicae*, regardless of infection status and spore concentration, our findings indicate that at specific concentrations, *B. bassiana* increases the survival and fecundity of *M. persicae*. This effect was also observed across a generation of *M. persicae*. Furthermore, we observed that *R. viridis* provides no clear protective effect against *B. bassiana*. These findings have broader implications for the effective application of fungal-based insecticides to control aphids while minimizing the occurrence of hormesis.

## RESULTS

### Aphid survival post-fungal exposure

The survival of *R. viridis*-free (R-) and *R. viridis*-infected (R+) *M. persicae* was monitored daily for 14 days following exposure to *B. bassiana* strain PRRI 5539 on leaf discs in Petri dishes with differing spore concentrations. We found significant effects of *B. bassiana* concentration on the survival of R-(Log-Rank test [LR]: χ2 = 557, df = 3, *p* < 0.001) and R+ (LR: χ2 = 434, df = 3, *p* < 0.001) aphids. Compared to the control group (0.1% Tween-80 solution), exposure to 1 × 10^4^ conidia mL^−1^ *B. bassiana* spores increased survival at day 14 from 11% to 56% for R+ (Figure 1a) and from 31% to 75% for R-(Figure 1b). However, exposure to 6 × 10^6^ conidia mL^−1^ and 1 × 10^8^ conidia mL^−1^ *B. bassiana* spores both decreased the survival of R+ and R-aphids. These findings provide evidence supporting the presence of hormesis in both the R- and R+ aphid populations, characterized by an increase in fitness following exposure to a toxic agent, which remains effective in eliminating the target at higher concentrations (Calabrese and Blain 2005).

**Figure 1.**
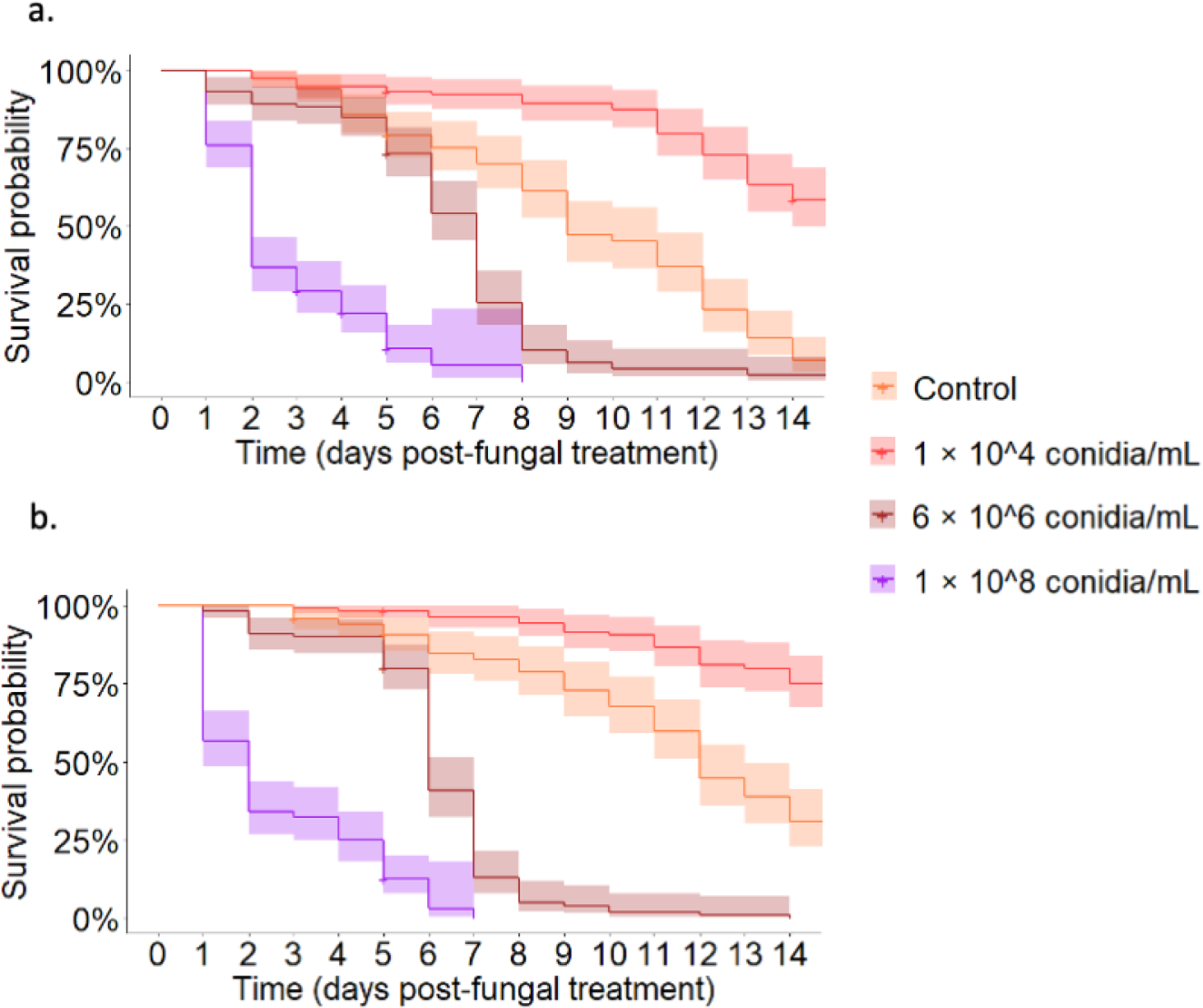
Kaplan-Meier curves showing the survival of (a) R+ and (b) R-*Myzus persicae* post-exposure to three different concentrations of *Beauveria bassiana* spores compared to the control. We tested *N* = 120 aphids per exposure group (8 replicates of 15 aphids per Petri dish). The shaded areas represent 95% confidence intervals. Log-rank tests (LR) and pairwise comparisons between treatments were performed to compare differences between survival curves with α = 0.05.

We observed a significant adverse effect of *R. viridis* on aphid survival in both the control group (R+ vs. R-aphids, LR χ2 = 21.8, df = 1, *p* < 0.001) and in the presence of 1 × 10^4^ conidia mL^−1^ of *B. bassiana* spores (R+ vs. R-aphids, LR χ2 = 6.7, df = 1, *p* < 0.009). On average, *R. viridis*-free aphids lived three days longer than *R. viridis*-infected aphids. Interestingly, this deleterious effect of *R. viridis* was not observed in aphids exposed to 6 × 10^6^ conidia mL^−1^ (R+ vs. R-aphids, LR χ2 = 2.2, df = 1, *p* = 0.100) and 1 × 10^8^ conidia mL^−1^ (R+ vs. R-aphids, χ2 = 0.5, df = 1, *p* = 0.500) of entomopathogenic fungi spores.

### Aphid fecundity post-fungal treatment

*Beaveria bassiana* exposure significantly influenced the fecundity of R- and R+ aphids (Type III ANOVA for Linear Mixed-Model [LMM]: F_3,56_ = 121.8771, *p* < 0.001). Exposure to entomopathogenic fungi spores with concentrations of 1 × 10^4^ conidia mL^−1^ and 6 × 10^6^ conidia mL^−1^, increased the fecundity of both R+ and R-aphids starting at day 2 post-fungal exposure, while exposure to 1 × 10^8^ conidia mL^−1^ entomopathogenic fungi spores significantly decreased the fecundity of both R- and R+ aphids (Figure 2a-b). *Rickettsiella viridis* also had a significant effect on fecundity (LMM: F_1,56_ = 24.01, *p* < 0.001), where R+ aphids had lower fecundity compared to R-aphids (Figure 2a-b). There was no significant interaction between *R. viridis* and *B. bassiana* spore concentration on fecundity (LMM: F_3,56_ = 1.93, *p* = 0.14).

**Figure 2.**
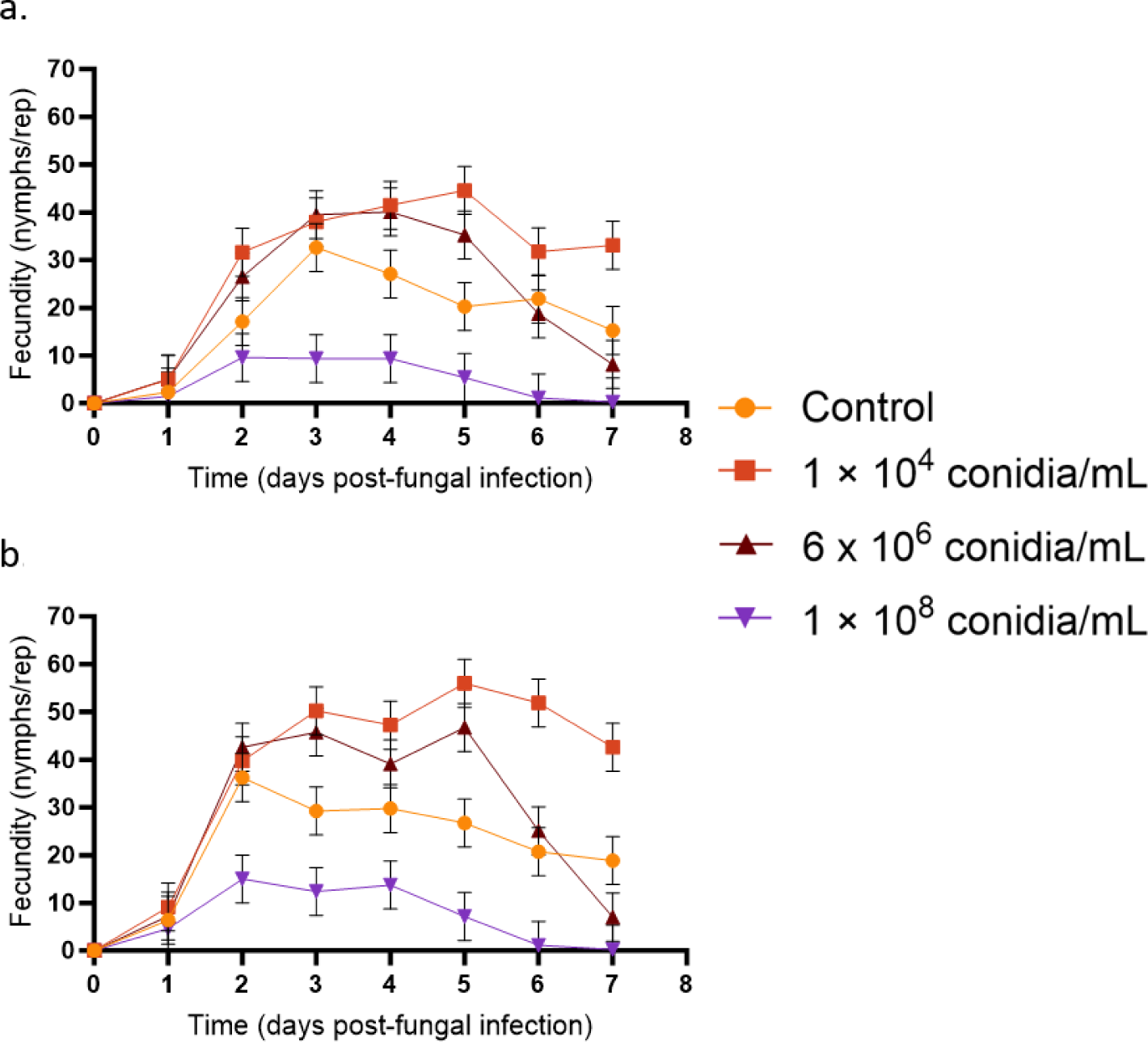
Fecundity of (a) R+ and (b) R-*Myzus persicae* over 7 days post-exposure to three different concentrations of *Beauveria bassiana* spores. We tested *N* = 120 aphids per exposure group (8 replicates of 15 aphids per Petri dish). Bars represent 95% confidence intervals. Overlapping bars indicate that the data points are not significantly different from one another.

### Aphid fecundity and development time in the second generation

In light of a robust hormetic response in aphids following exposure to *B. bassiana* spores, we explored the transgenerational persistence of hormesis by investigating the development time and fecundity of R+ and R-aphids exposed to *B. bassiana* in the previous generation. We found that *R. viridis* had no significant impact on the fecundity of F1 aphids (LMM: F_1, 122_ = 0.795, *p* = 0.374). On the other hand, we observed a significant effect of parental *B. bassiana* exposure on the fecundity of F1 aphids (LMM: F_3,122_ = 34.48, *p* < 0.001), where fecundity increased at all doses compared to control aphids in both R- and R+ populations (Figure 3a-b). Interestingly, the offspring of R+ and R-aphids exposed to 1 × 10^8^ conidia mL^−1^ produced more nymphs compared to other treatments. Furthermore, we observed a significant interaction between *R. viridis* infection and *B. bassiana* exposure on F1 aphid fecundity (LMM: F_3, 122_ = 34.47, *p* < 0.001), where the fecundity increase of *B. bassiana* exposure was more pronounced in the R+ population.

**Figure 3.**
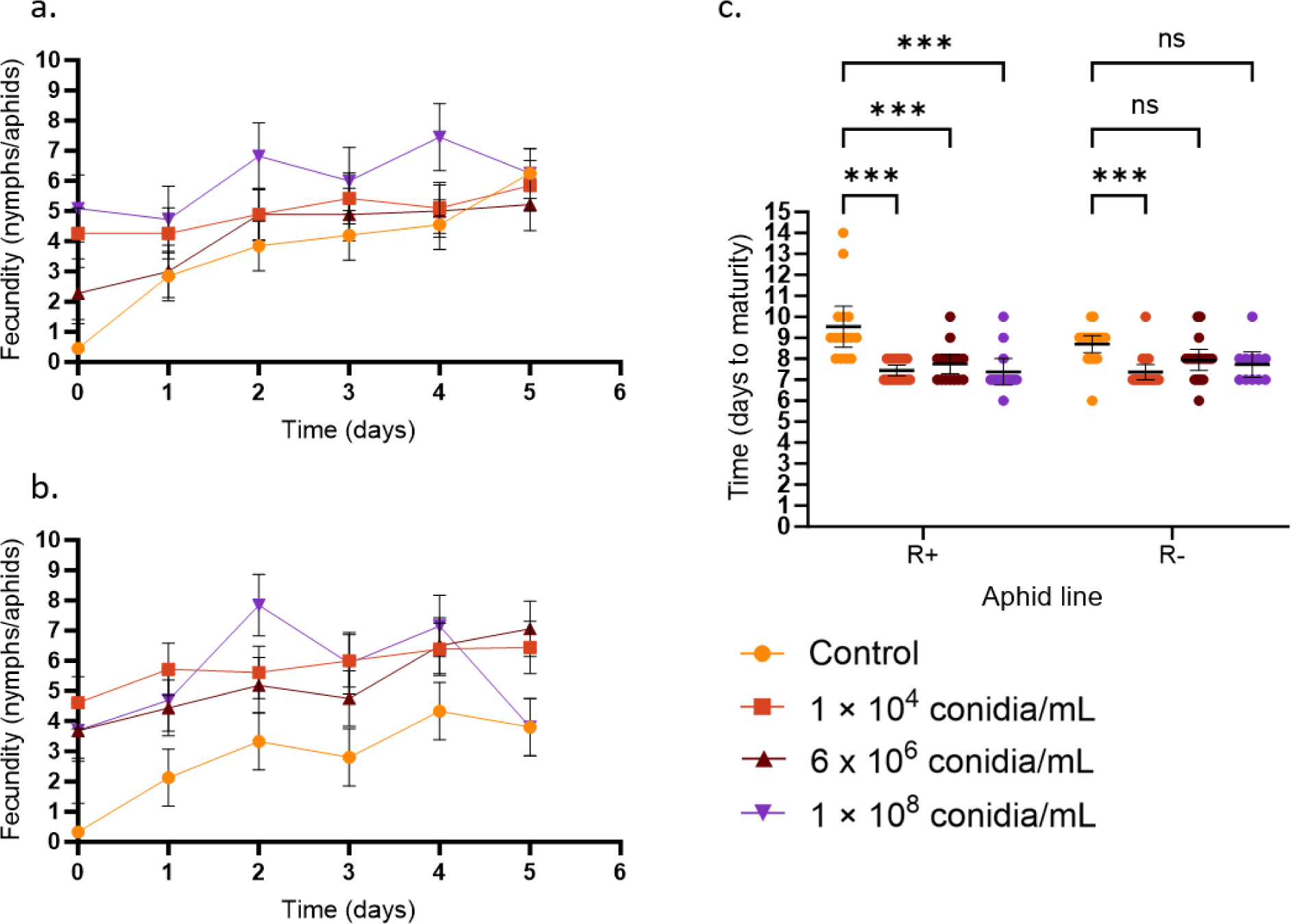
Fecundity of (a) R+ and (b) R-F1 aphids and (c) development time of R+ and R-F1 aphids following parental exposure to *Beauveria bassiana* spores. We tested 10-20 individual aphids per group. The start of fecundity measurement begun when aphids were 8 days old and some of the aphids have started reproducing. Bars represent 95% confidence intervals. Overlapping bars indicate that the data points are not significantly different from one another. ‘***’ represents = ‘p < 0.001’. Tukey’s honestly significant different (HSD) test was used to adjust *p*-values in Figure 3c. Only multiple comparisons that show a significant difference are shown.

We then assessed the development time of F1 aphids and found no significant influence of *R. viridis* infection (LM: F_1, 122_ = 0.27, *p* = 0.603). However, there was a significant impact of parental exposure to *B. bassiana* on the development time of F1 aphids (LM: F_3, 122_ = 20.42, *p* < 001). Both R+ and R-F1 aphids from mothers exposed to *B. bassiana* spores exhibited an accelerated time, approximately 1.5 days earlier than the control F1 R+ and R-aphids (Figure 3c). This effect was significant for all concentrations in the R+ aphid group, while in the R-aphid group, only F1 aphids from parents exposed to 1 × 10^4^ conidia mL^−1^ showed a significant difference from the control (Figure 3c). We did not find any significant interaction between parental *B. bassiana* exposure and *R. viridis* infection on the development time of F1 aphids (LM: F_3, 122_ = 2.16, *p* = 0.096). These findings, coupled with the F1 fecundity measurements, validate the enduring impact of hormesis across generations.

### Whole Plant Experiment with Different Aphid Clones

Following our initial leaf disc experiment, we replicated it using whole plants to investigate aphid hormesis at the low concentration of 1× 10^4^ conidia mL^−1^ *B. bassiana* spores. We included an additional *R. viridis*-free clonal line, denoted as R2-, alongside the R+ and R-populations. We found a significant positive impact of exposure to 1 × 10^4^ conidia mL^−1^ entomopathogenic fungi spores on aphid fecundity (LM: F_2, 41_ = 5.604, *p* = 0.023). In pairwise comparisons within lines, there was a significant increase in fecundity in the R-line, corroborating the hormetic response observed in the leaf disc experiment (Figure 4). However, while there was an increase in the fecundity of R+ aphids, this was not significant.

**Figure 4.**
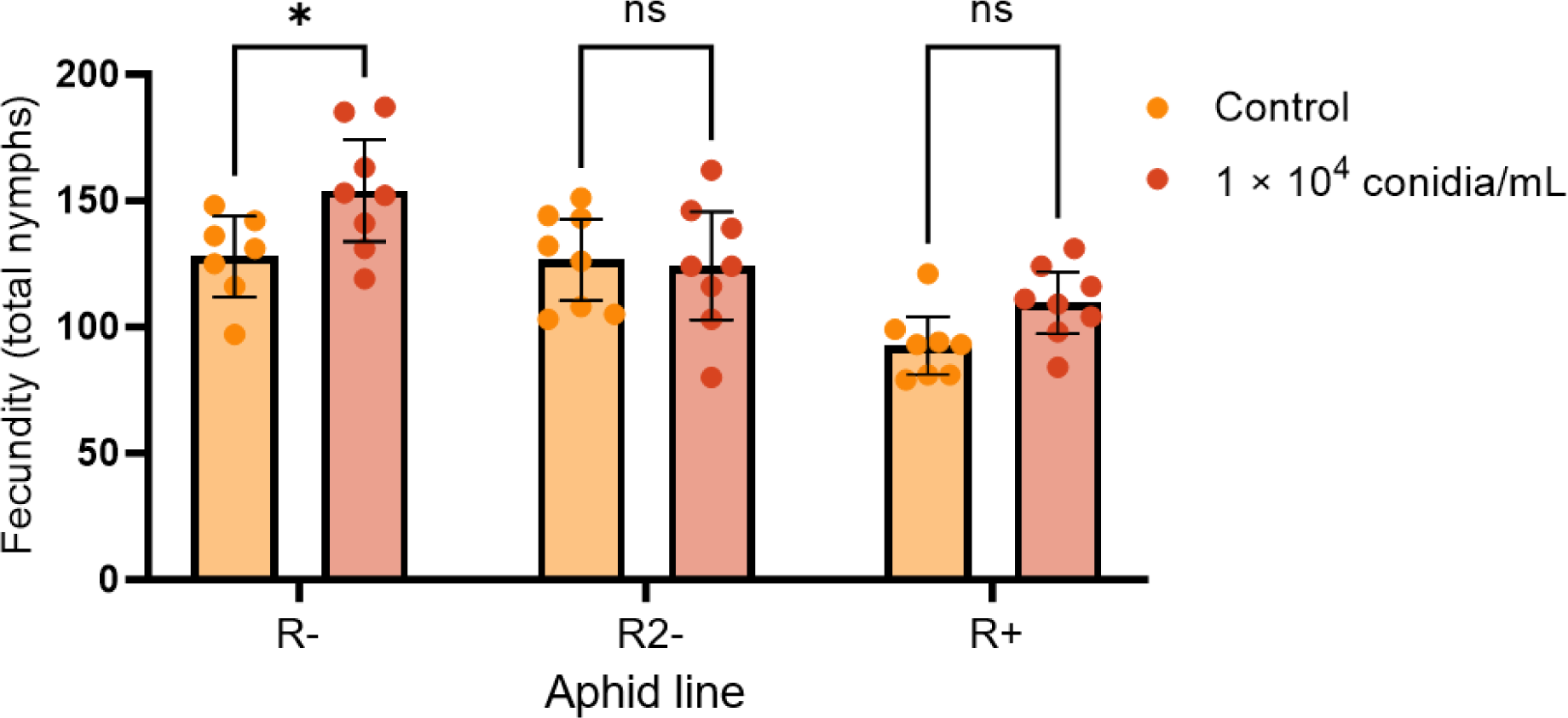
Fecundity of *Myzus persicae* on whole plants over a period of 7 days post-exposure to a low concentration of *Beauveria bassiana* spores. We measured fecundity from 5 adult female aphids per plant with 8 plants per line. Bars represent 95% confidence intervals. ‘*’ = ‘p < 0.05’. The Bonferroni method was used to adjust p-values. Dots represent the total fecundity from a single replicate plant.

In contrast, exposure to fungal spores had no discernible effect on the fecundity of R2-aphids, indicating differing responses to fungal exposure among aphid lines, including the exhibition of hormetic responses. Conversely, the most significant factor affecting fecundity was the aphid clonal lines itself (LM: F_2, 41_ = 16.14, *p* < 0.001). Specifically, as illustrated in Figure 4, *R. viridis*-infected aphid line (R+) exhibited lower fecundity on average compared to *R. viridis*-free aphids (R- and R2-), aligning with our previous findings that *R. viridis* has a detrimental impact on aphid fecundity. Furthermore, we did not observe any significant interaction between aphid clonal lines and treatment in relation to aphid fecundity (LM: F_2, 41_ = 2.11, *p* = 0.135).

## DISCUSSION

This study demonstrates that exposure to 1 × 10^4^ conidia mL^−1^ *B. bassiana* strain PRRI 5339 (entomopathogenic fungi) spores increase the survival and fecundity of *Myzus persicae*, regardless of *R. viridis* infection. Notably, these effects were also evident in the offspring of aphids exposed to *B. bassiana* regardless of the spore concentration. We also found an increase in fecundity in a whole-plant context for some clonal lines. Additionally, we confirmed that *R. viridis* had deleterious effects on aphid survival and fecundity but with a potential protective effect conferred against entomopathogenic fungi, particularly at higher concentrations of entomopathogenic fungi spores. The entomopathogenic fungi-induced increases in survival and fecundity in some *M. persicae* clonal lines highlights the necessity for further research to for developing entomopathogenic fungi-based bioinsecticide applications.

Hormesis is a phenomenon in which exposure to low, often sublethal concentrations of a toxic agent increases the fitness of the exposed organism, while at higher concentrations, the agent’s effects remain lethal (Calabrese 2014). In our study, exposure to entomopathogenic fungi spores with a concentration of 1 × 10^4^ conidia mL^−1^ increased both fecundity and survival of aphids, while exposure to 1 × 10^8^ conidia mL^−1^ caused rapid mortality and greatly reduced fecundity, strongly indicating hormesis. While reports on chemical-based insecticide-induced hormesis in aphids are plentiful (Chen et al. 2020; Li et al. 2023), reports on biopesticides are limited, with one study finding that exposure to endophytic *B. bassiana* increased the fecundity of black bean aphids (*Aphis fabae* Scopoli) and this effect persisted across two aphid generations (Jensen et al., 2019).

We also found a transgenerational effect of *B. bassiana* exposure, where hormesis occurred in both *R. viridis*-free and *R. viridis*-infected F1 *M. persicae*. Previous studies have observed transgenerational hormetic responses when aphids are continuously exposed to low concentrations of a toxic agent (Ayyanath et al. 2013). Interestingly, in our study hormesis also was observed under *B. bassiana*-free conditions, where F1 aphid nymphs were born on fresh leaf discs without *B. bassiana* and subsequently isolated from their entomopathogenic fungi-infected mothers. We hypothesize this hormesis was the result of either residual *B. bassiana* spores or transgenerational selection that favored increased fecundity. The increase in F1 fecundity was largest in the progeny of aphids exposed to 1 × 10^8^ conidia mL^−1^ in support of a selection hypothesis.

The mechanism underlying the adaptive hormetic response remains elusive. Jensen and colleagues (2019) argue that elevated-plant defense mechanisms due to endophytic entomopathogenic fungi, stresses the aphids and leads to hormesis. In this study, we suggest that *B. bassiana* may also directly stress the aphids. Two factors place the aphids under constant risk of mortality. First, *B. bassiana* produces secondary metabolites with immunosuppressive and insecticidal properties, such as beauvericin, bassianin, bassianolide, beauverolides, and others (Wang et al. 2021). Second, *B. bassiana* in low spore concentrations establishes long-term infections without eliminating the aphids (Dubovskiy et al. 2013). In response to the constant threat, aphids may invest in increased fecundity (Barribeau et al. 2010).

Hormetic effects induced by entomopathogenic fungi such as *B. bassiana* raises concerns for potential field applications, where some areas receive lower doses of the pesticide. With hormesis, this could potentially lead to a quick resurgence of pest populations that are well adapted to the pesticides (Guedes and Cutler 2014). Future work is required to predict the likelihood and extent of hormesis in a field context, which will likely to depend on geographic location and resistance levels given that only some clonal lines exhibited increased fecundity in our whole plant experiments. This finding aligns with the previous literature where clonal variation influences aphid resistance to pathogens (Ferrari et al. 2001).

Some of our data are consistent with a protective role of *R. viridis* in the novel host *M. persicae*. Prior literature suggests that *R. viridis* can protect *A. pisum* from *Pandora neoaphidis* (Łukasik, van Asch, et al. 2013). Here we found modest increases in the survival of *R. viridis*-infected *M. persicae* post-exposure to high concentrations of *B. bassiana* compared to *R. viridis*-free *M. persicae*, despite *R. viridis* decreasing survival in the control. Furthermore, in the offspring of parents exposed to *B. bassiana*, *R. viridis*-infected *M. persicae* tended to have higher fecundity than *R. viridis*-free *M. persicae*. There are currently no described mechanisms underlying the protective effects conferred by endosymbionts against entomopathogenic fungi, but could involve combinations of these factors: (1) endosymbiont-induced “immune priming”, where endosymbiont infection increases the production of antimicrobial peptides (Eleftherianos et al. 2013); (2) immune adaptation, where host resistance to fungal pathogen gradually increased (Dubovskiy et al. 2013); (3) endosymbionts facilitate detoxification, in this case detoxification of *B. bassiana* toxins such as beauvericin (Blanton and Peterson 2020); (4) endosymbionts produce antifungal compounds (Kett et al. 2021); (5) endosymbionts augment formation of host cuticle (Kanyile et al. 2022); (6) general nutrition provisioning to support host neutralizing active fungal infection (Brownlie et al. 2009; Hosokawa et al. 2010). It might also be caused by selection imposed by *B. bassiana* across generations. This present study demonstrates that the protective role of *R. viridis* against *B. bassiana* is limited, and fungal-based pesticides, especially high concentrations *Beauveria bassiana* (*i.e.,* 1 × 10^8^ conidia mL^−1^) is still effective in controlling the aphids including *R. viridis*-infected *M. persicae*. Moreover, in the context of entomopathogenic fungi -induced or chemical-induced hormesis, *R. viridis* may have a significant role in mitigating the hormetic response.

In summary, our study has revealed that a specific clonal line of *M. persicae* exhibits hormesis in response to *B. bassiana* infection and the beneficial adaptive response extends to the second generation. This is also evident on whole plants treated with *B. bassiana*. Our study is limited to only testing one species and pathogen which need more research to show the hormesis effects among different species and endosymbionts. This hormetic response raises concerns for field applications but further research is required to monitor its prevalence under realistic conditions and to understand its underlying mechanisms.

## METHODS AND MATERIALS

### Aphid rearing

*The M. persicae* line harbouring *R. viridis* (R+) was generated with microinjection from the native donor *A. pisum* (Gu et al. 2023) using a single clonal line, GPA_V_20191007_08 as the recipient. The *R. viridis*-free line (R-) was from the same clone, and was originally collected in Queensland, Australia from canola (*Brassica napus*). The R+ line has been maintained in the lab for more than 60 generations with a 100% transmission rate. Prior to experiments, 30-40 adults per line (R+ and R-) were on placed on whole bok choy (*Brassica rapa* subsp. chinensis) plants aged approximately 9 weeks in 30 × 30 × 62 cm cages (160 µm aperture mesh) to grow for approximately 2.5 weeks. These aphids were kept in climate controlled rooms at 19 – 21 °C with a 16:8 light and dark cycle. At the end of population expansion period, 20 – 30 adult apterous aphids were transferred onto bok choy leaves on 1% agar in 100 mm Petri dishes. Twenty-four hours after the transfer, nymphs were counted and 20 – 40 nymphs placed on fresh leaves in new 100 mm Petri dishes to avoid stressing the aphids. The nymphs were reared for 7-8 days in an incubator at 19 °C with a 16:8 light and dark cycle. During the rearing period, the leaves were changed once on day 5. Another clonal line of *R. viridis*-free *M. persicae* GPA_Q_20211116_26 (designated as R2-, collected in Queensland, Australia from sugarloaf cabbage [*Brassica oleracea* var. capitata]) was set up for the whole plant experiments following the same procedure as described above.

### Leaf dip bioassays and survival and fecundity measurements

A stock suspension containing *Beauveria bassiana* strain PRRI 5539 spores (1.7 × 10^9^ spores mL^−1^) was obtained from BASF^®^. Three spore concentrations (i.e., 1 × 10^8^, 6 × 10^6^, and 1 × 10^4^ spores mL^−1^) were prepared by diluting the stock suspension using the appropriate amount of 0.1% Tween-80 solution (Akmal et al. 2013; Sayed et al. 2021). To test the concentration of *B. bassiana* spores, the stock solution concentration was determined using a haemocytometer and a light microscope at 400 times magnification. Prior to assays, bok choy leaves were sterilised using 0.2% bleach and 0.1% Tween-80 solution and followed by a 5-10 second 100% ethanol wash. Leaves were then washed three times with dH_2_O to remove residual ethanol and bleach. A leaf bioassay procedure was performed, where bok choy (*Brassica rapa* subsp. chinensis) leaves were cut into 35 mm discs which were submerged in each prepared spore concentration for 5 minutes and constantly agitated by gently swirling the container to prevent clumping of spores. For the control, leaf discs were dipped in a 0.1% Tween-80 solution for 5 minutes. Leaves were left to dry in a fume hood then placed on 1% agar in 60 mm petri dishes by gently pressing against the agar (1 leaf disc per petri dish). Each treatment group was replicated eight times and each replicate contained 15 aphids aged 7-8 days (4^th^ instar nymph). Aphids were monitored daily for 14 days post fungal exposure. Petri dishes were stored inside a climate controlled incubator (19 °C with a 16:8 light and dark cycle). Aphids were exposed for the duration of three days and subsequently, fresh Bok choy leaves were changed every three days after the exposure. Aphid fecundity was monitored, and nymph removal was performed daily. Aphid survival was assessed based on aphids body colour and aphids remained active when probed with a soft brush each day.

### Transgenerational effects on development time and fecundity

On day 5 after fungal exposure, the offspring (F1: 1 day old; 1^st^ instar) of aphids exposed to each concentration of *B. bassiana* were haphazardly selected and placed individually on fresh 25 mm bok choy leaf discs in a 35 mm petri dish containing 1% agar. Ten to twenty replicates per concentration for each line were set up. Development time was monitored daily by determining the time for each aphid to produce its first nymph. Fecundity was scored by counting and removing F2 nymphs daily for 14 days since the start of the experiment. The petri dishes were kept at 19 °C with a 16:8 light and dark cycle.

### Whole plant experiment

One month old bok choy plants were used and separated into two treatments, one with no *B. bassiana* exposure and the other exposed to a low concentration of *B. bassiana* spores (10^4^ conidia/mL diluted in 0.1% Tween-80 solution) using a 1L hand-spray. Three sprays were applied at each of the five different positions with a distance approximately 20 - 30 cm: south, north, west, and east relative to the centre of the container, as well as from the top side. Three lines of aphids R-, R+, and R2-were transferred onto each plant (5 adult female apterous aphids/plant). Each treatment group was replicated 8 times. The bok choy plants were covered with clear perforated plastic bag and maintained 19 °C with a 16:8 light and dark cycle by watering every 2 days. The fecundity of aphids was scored by counting and removing nymphs daily for 7 days.

### Aphids DNA extraction and quantitative PCR

From each *B. bassiana* spore exposed and control group, one to two surviving adult aphids per replicate were collected on day 5 post fungal exposure. Aphid DNA was extracted using 150 μL of 5% Chelex^®^ 100 resin (Bio-Rad Laboratories, Hercules, CA). Aphid tissue was homogenized using a Qiagen TissueLyser II^®^ with 2-3 mm glass beads (AjaxFineChem, Taren Point, NSW, Australia; Cat. No. 1700-500G) at an oscillation frequency of 25 Hz for 1 minute. Subsequently, 2 μL of Proteinase K (20 mg mL-1) was added to the homogenates. The homogenates were incubated at 65°C for 30 minutes, followed by a 10-minute incubation at 90°C. After centrifugation for 4 minutes at 20,800 × g (Eppendorf Centrifuge 5417 C), aphid genomic DNA in the supernatant was diluted in molecular-grade H_2_O with a 3-fold dilution factor.

A quantitative polymerase chain reaction (qPCR) assay was performed to detect and quantify the abundance of *B. aphidicola* and *R. viridis*, relative to the actin gene (Lee et al. 2012, Gu et al. 2023) with LightCycler^®^ 480 High Resolution Melting Master (HRMM) kit (Roche; Cat. No. 04909631001, Roche Diagnostics Australia Pty. Ltd., Castle Hill New South Wales, Australia) and Immolase™ DNA polymerase (5 units µL^−1^) (Bioline; Cat. No. BIO-21047). We included 4 wells of aphid DNA extracts with confirmed *R. viridis* infection statuses (2 R+ and 2 R-) and 2 wells with only primers as controls in each qPCR run. We used three primer pairs: (1) Universal aphid actin (forward: 5’GTGATGGTGTATCTCACACTGTC; reverse: 5’AGCAGTGGTGGTGAAACTG); (2) *B. aphidicola* 16s rRNA (forward: 5’AAAGCTTGCTTTCTTGTCG; reverse: 5’GGGTTCATCCAAAAGCATG); (3) *R. viridis* 16s rRNA (forward: 5’GGGCCTTGCGCTCTAGGT; reverse: 5’TGGGTACCGTCACAGTAATCGA). We performed two technical replicates per individual aphid. The densities of *R. viridis* and *B. aphidicola* were normalised against the internal gene expression (i.e., aphid actin gene). The average cycle threshold (CT) values of both *R. viridis* and *B. aphidicola* were subtracted with the average CT value of the aphid actin. The unit of abundance represented in the figures are log transformed log_10_ (2^ΔCT^).

### Statistical Analysis

All statistical analyses were performed using R and graphical representations of the data were created using both GraphPad Prism version 10.1.10 and R. Survival of *M. persicae* was analysed using the *survival* package (version 3.5-5) (Therneau et al. 2023), which contains log-rank test function (LR) to investigate the overall differences between survival curves. Kaplan-Meier curves (the survival of *M. persicae* over the period of observation) were created using the *ggplot2* (version 3.4.2) and *survminer* packages (version 0.4.9) (Kassambara et al. 2021; Wickham et al. 2023).

The fecundity of F0 and F1 *M. persicae* was analysed using linear mixed models (LMMs) where fecundity was set as the response variable where *R. viridis* infection, *B. bassiana* exposure spore concentrations, and time were set as fixed effects, and biological replicates was set as the random effect. F0 and F1 *M. persicae* fecundity were considered to be normally distributed. The R-studio packages used for these LMMs analysis and visualizations were as follows: *lme4* (version 1.1-33), *multcomp* (version 1.4-23), and *ggplot2* (version 3.4.2), DHARMa, MuMIn (Bates et al. 2023; Hothorn et al. 2023; Patil and Powell 2023; Bartoń 2023; Hartig and Lohse 2022). Type-III analysis of variance (ANOVA) was used to assess the overall effect of the variables explained by LMMs and post-hoc test of LMMs was performed using *emmeans* (version 1.8.5) (Lenth et al. 2023) with Bonferroni method used to adjust the *p*-values. The significance threshold value was set to α = 0.05.

The development time of F1 *M. persicae* was analysed using two-way analysis of variance (ANOVA) where the response variable was time which was defined as days to first nymphs, and the effect variables were *Rickettsiella viridis* infection and *Beauveria bassiana* exposure spore concentration. Development time was considered to follow a normal distribution.

Two-way ANOVA was also used to investigate the differences between the density (log_10_ (2^ΔCT^)) of *Buchnera aphidicola* and *Rickettsiella viridis* in F0 *M. persicae* that were exposed to three different concentrations of *B. bassiana*. The response variable was the endosymbiont density (log_10_ (2^ΔCT^)), and the effect variables were combination of *R. viridis* infection and *B. bassiana* spore concentrations for investigating *B. aphidicola* density and *B. bassiana* spore concentrations for investigating *R. viridis* density.

The whole plant experiments were also statistically analysed using two-way ANOVA. For these two-way ANOVA analyses Tukey’s Honestly Significant Difference method was used as the post-hoc test. The significance threshold value was also set to α = 0.05. GraphPad Prism version 10.1.10 were used for two-way ANOVA analyses and to create most of the figures shown in this study.

## ACKNOWLEDGEMENTS

We would like to thank Alex Gill, Sonia Sharma, Nick Bell, Joshua Thia, Nancy Endersby-Harshman, Qiong Yang, Ash Dorai, Eloise Ansermin, Zhenyu Zhang, Ashley Callahan, and other members of PEARG for their support, insight, intellectual discussions, and technical assistance.

