## Supplemental figure 1 for "Hormetic effect induced by *Beauveria bassiana* in *Myzus persicae*"

**APPENDIX**

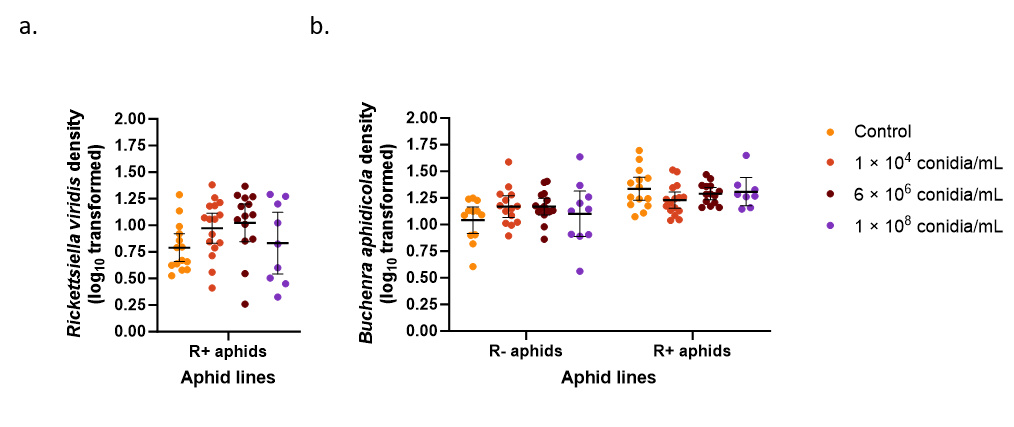

***Supplementary Figure 1.*** Abundance of F0 *M. persicae* endosymbiotic bacteria. We measured the density of *Rickettsiella viridis* (a) and *Buchnera aphidicola* (b) across different concentrations of *Beauveria bassiana* (1 × 10^4^, 6 × 10^6^, 1 × 10^8^ conidia mL^-1^). Lines and error bars represent means and 95% confidence intervals.
